# The heat shock protein 70 (Hsp70) family of chaperones is essential for rhinovirus replication

**DOI:** 10.64898/2026.08.13.742791

**Authors:** Matthew T. James, Courtney Dane, Alanna O. Moore, Aurélie Mousnier

## Abstract

2.

Rhinoviruses (RVs) are the predominant cause of the common cold and a major trigger of acute asthma exacerbations. Yet, unlike related enteroviruses such as poliovirus (PV) and enterovirus A71 (EV-A71), no approved vaccines or antivirals exist. Because enteroviruses depend heavily on host factors for replication, cellular proteins that support the replication of multiple enteroviruses have emerged as attractive broad-spectrum antiviral targets that may offer a higher barrier to resistance than virus-targeted therapies. The 70-kDa heat shock protein (Hsp70) family, a highly conserved class of molecular chaperones, is required for the replication of several enteroviruses, including EV-A71 and coxsackievirus A16, but whether RVs share this dependency was unknown.

Here, we show that two mechanistically distinct small-molecule inhibitors of the Hsp70 family abolish replication of RV-A16, a clinically relevant RV type widely used in asthma research. Using siRNA knockdown, we demonstrate a role for HSPA8, the major constitutively expressed Hsp70 isoform, in RV replication. We further demonstrate that Hsp70 activity is indispensable for viral translation, identifying this as a key Hsp70-dependent step in the RV replication cycle. Together, these findings establish Hsp70 chaperones as essential host factors for RV replication and strengthen the rationale for targeting them as a broad-spectrum antiviral strategy.

**Impact statement:** Rhinoviruses are the predominant cause of the common cold and major triggers of exacerbations in people with asthma and chronic obstructive pulmonary disease, yet no licensed antiviral therapies exist. Although several enteroviruses are known to require Hsp70 chaperones for replication, whether rhinoviruses share this dependence was unknown. Here, we address this gap by demonstrating that Hsp70 chaperones are essential host factors for rhinovirus replication and are required for viral translation. These findings advance our understanding of how rhinoviruses exploit the host cell machinery and broaden the evidence supporting Hsp70 chaperones as potential antiviral targets across the *Enterovirus* genus. As host-targeted therapies may be less vulnerable to resistance than direct-acting antivirals, these findings represent an important step towards the development of urgently needed anti-rhinoviral therapeutics and will be of interest to virologists, respiratory clinicians and antiviral drug developers.

## 4. Introduction

Rhinoviruses (RVs) are among the most pervasive viral pathogens of humans, responsible for millions of infections annually and the most frequently detected cause of acute respiratory illness worldwide^1^. RVs predominantly infect the upper respiratory tract, causing common colds, which can lead to complications such as rhinosinusitis and otitis media^2,3^. Although these infections are typically self-limiting, their high frequency - most individuals experience multiple episodes each year - imposes substantial health and socioeconomic burdens, including lost productivity, healthcare utilisation, and inappropriate antibiotic use, which may contribute to antibiotic resistance^4,5^.

RV infections are also a significant cause of lower respiratory tract disease. RVs are indeed frequently detected as the sole respiratory pathogen in hospitalised patients and rank among the most commonly identified viruses in severe respiratory infections^6,7^. RVs are the dominant trigger for asthma exacerbations, detected in over half of cases, many of which require emergency care or hospitalisation and may occasionally prove fatal^8,9^. RVs are also frequently identified during acute exacerbations of chronic obstructive pulmonary disease (COPD), a leading cause of hospitalisation and mortality worldwide^10^.

Despite their prevalence and clinical impact, there are no licensed vaccines or antivirals against RVs. A principal barrier to controlling RV disease is their extensive genetic diversity. RVs comprise three species (commonly named RV-A, -B and -C) and over 170 antigenically distinct (geno)types that circulate globally and year-round^11^, with minimal cross-protective immunity^12^. Nevertheless, genetically diverse RVs often depend on a common set of host proteins to replicate. Some of these proteins, such as GBF1^13–19^, PI4KIIIβ^13,20–23^, and SETD3^24^, are indeed essential for the replication of multiple RVs and related enteroviruses such as polioviruses (PVs) and enterovirus A71 (EV-A71). Identifying these “pan-enterovirus” host dependencies and defining their role in viral replication may enable the development of broad-spectrum antiviral strategies targeting them. In addition, these strategies may be less easily circumvented by viral genetic variation than strategies that directly target the virus.

Cellular protein homeostasis is maintained by molecular chaperones, which are co-opted by viruses to facilitate their replication. Among these, 70-kDa heat shock proteins (Hsp70s) have been shown to support the replication of several RNA viruses, including some enteroviruses^25–33^. The Hsp70 family is highly conserved and ubiquitously expressed in eukaryotic cells. Together with co-chaperones, Hsp70s maintain cellular protein homeostasis through diverse functions, including the folding of newly synthesised proteins^34^, the assembly and disassembly of protein complexes^35,36^, the import of proteins into organelles^37,38^, and the disaggregation of protein aggregates^39^. The human genome encodes 13 Hsp70 homologues with distinct subcellular localisations and expression profiles. Some, such as HSPA8 (also known as Hsc70), are constitutively expressed, whereas others, including HSPA1A and HSPA1B, are stress-inducible. Hsp70 proteins bind to short hydrophobic peptide sequences exposed in unfolded or misfolded substrates. Their chaperone activity is driven by cycles of substrate binding and release, regulated by ATP binding and hydrolysis and two classes of co-chaperones: J-domain proteins (JDPs, also known as DnaJs or Hsp40s) and nucleotide exchange factors (NEFs). JDPs help deliver substrate proteins to Hsp70 and stimulate ATP hydrolysis, stabilising substrate binding, while NEFs exchange ADP for ATP on Hsp70, enabling substrate release^35^.

Members of the Hsp70 family have been implicated in the replication of several enteroviruses^26,31,32,40^. Notably, Su *et al.* demonstrated that pan-Hsp70 inhibition with the small molecule JG40 - an allosteric inhibitor that blocks Hsp70-NEF interactions - inhibits multiple stages of the EV-A71 replication cycle, including internal ribosomal entry site (IRES)-mediated viral RNA translation^31^. JG40 also inhibited the replication of several other enteroviruses, including coxsackievirus A16 (CVA16), CVB1, CVB3, and echovirus 11 (E11)^31^. However, despite these findings, direct evidence supporting a role for Hsp70 chaperones in RV replication is lacking.

In this study, we investigated the roles of Hsp70 chaperones in the replication cycle of RV-A16, a clinically relevant RV type widely used as a model in asthma studies^41^. We demonstrate that two mechanistically distinct small molecule inhibitors of all Hsp70s completely block RV-A16 replication, indicating an essential requirement for their activity. Using siRNA-mediated knockdown, we further show that the major constitutively expressed Hsp70 family member, HSPA8, as well as two JDP co-chaperones, specifically contribute to RV-A16 replication. We further demonstrate that Hsp70 chaperones are indispensable for IRES-mediated translation of the viral RNA, providing mechanistic insight into their role in the viral replication cycle. This study expands our understanding of how RVs hijack host factors for replication, provides further evidence supporting Hsp70s as important “pan-enterovirus” host factors, and may open promising new avenues for antiviral development.

## 5. Methods

A detailed list of key resources used in this study is provided in Supplementary Table 1.

### Cells and viruses

Human cervical carcinoma HeLa-H1 cells (ATCC) were cultured as monolayers in complete medium (composition in Supplementary Table 1) at 37 °C, 5% CO2.

Rhinovirus A-16 (RV-A16; ATCC) was propagated in HeLa-H1 cells. Experimental virus stocks were generated by diluting master stock in infection medium (composition in Supplementary Table 1) and infecting two T175 flasks of HeLa-H1 cells (MOI=3), incubated at 33 °C, 5% CO_2_ until full cytopathic effect (CPE). All RV infections in this study were performed at 33 °C to replicate the upper respiratory tract^42^. After three freeze–thaw cycles, lysates were clarified (2,300 × g, 30 min, 4 °C), diluted 1:10 in infection medium, and twenty T175 flasks were infected and incubated as above. Following three freeze–thaw cycles and clarification, supernatants were aliquoted and stored at −80 °C, and one aliquot titrated by 50% tissue culture infectious dose (TCID_₅₀_) assay (see below).

### Virus titration

Experimental samples (cells and infection medium) were frozen at −80 °C at the indicated time points. After three freeze–thaw cycles, samples were harvested by scraping and the whole cell lysate collected. Experimental samples and freeze-thawed virus stock aliquots were clarified by centrifugation.

Clarified supernatants were serially diluted in titration medium (composition in Supplementary Table 1) in 96-well plates (4-6 technical replicates per sample), with eight 10-fold dilutions (10^-1^-10^-8^) prepared per replicate. HeLa-H1 cells (1×10^5^ cells/ml) were then seeded on top, and plates incubated at 33 °C, 5% CO_2_ for 5 days. CPE was scored by light microscopy and the TCID_50_/ml calculated by the Reed-Muench method^43^.

### RV infection of cell lines and inhibitor treatment

HeLa-H1 cells were infected at 95-100% confluence at the indicated multiplicity of infection (MOI), calculated from viral titres (TCID_50_/ml was converted to PFU/ml assuming 1 TCID_50_ = 0.7 PFU^44^; cell numbers were extrapolated from surface area assuming ≈13×10^6^ HeLa cells per confluent 75 cm^2^ flask). Virus was diluted in infection medium, adsorbed for 1 h at 33 °C, after which cells were washed five times and incubated in fresh infection medium at 33 °C, 5% CO_2_ until the indicated time point (for 0 h time points, cells were processed immediately after the 1 h adsorption).

For inhibitor studies, HeLa-H1 cells were treated with the indicated inhibitor (VER-155008, Neo Biotech; or pifithrin-µ, Merck), or dimethyl sulfoxide (DMSO). Cells were pre-treated prior to infection where indicated, and were re-treated during and after virus adsorption. At the indicated time points, cells were processed for viral titration, Western blotting or RT-qPCR (see respective sections).

For dose-effect assays, cells infected with RV-A16 (MOI=20) were treated with increasing concentrations of VER-155008 or DMSO and titrated at 0 h (input) and 6 h. For each experiment, technical replicates for VER-155008-treated conditions at 6 h were plotted as individual points. Technical replicates for DMSO/input conditions were averaged across all independent experiments and plotted as lines. For time-of-addition assays, cells infected with RV-A16 (MOI=20) were treated with 50 µM VER-155008 or DMSO at defined time points relative to the 0 h time point, and titrated at 6 h. One technical replicate was performed per condition per experiment.

### Cell viability assays

Cell viability was assessed using a resazurin-based assay in parallel to the corresponding infection or transfection experiments, with matched treatments and durations. HeLa-H1 cells were seeded in 100 µl complete medium in black, opaque 96-well plates with transparent bottoms (Greiner Bio-One). At 95-100% confluence, cells were treated with the inhibitor concentration(s) used in the corresponding experiment, or DMSO, in complete or infection medium, and incubated alongside the infection or transfection plate at 37 °C or 33 °C, 5% CO2 for the matched duration. At the endpoint, complete or infection medium alone was added to 5-6 blank wells. Resazurin working solution (Fluorochem; 0.15 mg/ml in PBS) was added (20 µl/well) and plates incubated 1 h at 37 °C, 5% CO2. Fluorescence was recorded (560 nm excitation / 590 nm emission; FLUOstar® Omega, BMG Labtech) and normalised to the mean of the blanks. Per experiment, 3-6 technical replicates were performed for drug-treated conditions and 5-6 each for DMSO and blanks.

### Cell transfections

For gene silencing using ON-TARGETplus SMARTpool siRNAs (4 siRNAs/gene; Dharmacon, see Supplementary Table 1), siRNA pools targeting HSPA8, DNAJA1 or DNAJC7, or a non-targeting control, were reverse-transfected with Lipofectamine® RNAiMAX (Thermo Fisher Scientific): per 96-well plate well, 4 pmol siRNA pool was diluted in 20 µl Opti-MEM® I (Thermo Fisher Scientific), 0.3 µl RNAiMAX added, and the mix incubated 30 min at RT, before 80 µl HeLa-H1 cells in complete medium were seeded on top (40 nM final). Cells were incubated 72 h to 95-100% confluence before infection with RV-A16 (MOI=20) and titration at 6 h. Per experiment, one technical replicate was performed per gene-targeting pool and six for the non-targeting control; each pool was expressed as a percentage of the mean of the non-targeting control across 3-4 experiments (mean of means). The corresponding viability assay was performed in parallel by resazurin assay, with three technical replicates per pool and six each for the control and blanks.

Single siRNAs (Eurofins, see Supplementary Table 1) targeting HSPA8^45^ or firefly luciferase were forward-transfected into HeLa-H1 cells (seeded to 30% confluence) with Lipofectamine® RNAiMAX: per 24-well plate well, 5 pmol siRNA in 50 µl Opti-MEM® I and 0.5 µl RNAiMAX in 50 µl Opti-MEM® I were each incubated 5 min at RT, combined and incubated 20 min, and added (100 µl) to cells in 400 µl low serum medium (10 nM final). Plates were incubated 48 h, after which low serum medium was replaced with complete medium for a further 24 h before processing or infection with RV-A16 (MOI=20). One technical replicate was performed per condition.

*In vitro* transcribed RNA was reverse-transfected using TransIT-mRNA and mRNA Boost reagents (Mirus Bio): RNA was diluted in Opti-MEM® I, the reagents added, and the mix incubated exactly 5 min at RT, before HeLa-H1 cells in complete medium were seeded on top (to reach 95-100% confluence the next day); amounts are given in Supplementary Table 2. Cells were incubated at 37 °C, 5% CO_2_ and processed at the indicated time points. One technical replicate was performed per condition per experiment.

### Viral RNA transfection assay

To assess whether Hsp70 activity is required at a step following viral entry, full-length RV-A16 RNA was transfected directly into cells, bypassing receptor-mediated entry and uncoating. The plasmid encoding full-length RV-A16 (pR16.11; a gift from Stanley Lemon and Kevin McKnight, University of North Carolina)^46^ was linearised and viral RNA *in vitro* transcribed and purified as described below (see ‘*In-vitro transcription and RNA purification’*). HeLa-H1 cells were pre-treated with 50 µM VER-155008 or DMSO for 1 h at 37 °C, then reverse-transfected with the transcribed RNA (see ‘Cell transfections’) in 24-well plates, with VER-155008 or DMSO maintained. At 8 h, cell lysates were processed for viral titration. Two technical replicates per condition were averaged per experiment.

### Nanoluciferase replicon assay

The replication-defective RV-A16 nanoluciferase replicon (pRVA-16-NL-YGAA), in which the 3D polymerase active-site motif YGDD is mutated to YGAA, was generated by site-directed mutagenesis (see ‘Plasmid construction’) and transcribed *in vitro* (see ‘In vitro transcription and RNA purification’). HeLa-H1 cells were pre-treated with 50 µM VER-155008, 178 µM cycloheximide (CHX; Thermo Fisher Scientific) or DMSO for 1 h at 37 °C, then transfected with the replicon RNA by reverse transfection, with the corresponding treatment maintained, in separate 96-well plates per time point (0 and 8 h). The 0 h plate was processed immediately; the 8 h plate was incubated at 37 °C, 5% CO_2_. Wells were processed using the Nano-Glo® Luciferase Assay (Promega) per instructions. 100 µl from each well was transferred to a fresh well in a white 96-well plate, and luminescence was recorded using the top lens optic with 1.0 s integration and auto-gain (FLUOstar® Omega, BMG Labtech). Per condition, 0 h luminescence was subtracted and values normalised to the 0 h-subtracted DMSO value at 8 h. Five technical replicates were averaged per condition per time.

### SDS-PAGE and Western Blotting

At the indicated time points, medium was removed and cells washed once in PBS. 2× Laemmli sample buffer (composition in Supplementary Table 1) supplemented with 0.2 M DTT (Fluorochem) was warmed, added per well or dish, and cells harvested with a scraper. Samples were vortexed, centrifuged (15,000 × g, 5 min, RT) and incubated at 100 °C, repeated until non-viscous. Samples were quantified using the dsDNA setting on a NanoDrop™ One spectrophotometer, normalised to equivalent DNA, and topped up to equal volumes with 2× Laemmli buffer. Bromophenol blue (Alfa Aesar) was added, and samples vortexed, incubated at 100 °C and centrifuged (15,000 × g, 5 min, RT) immediately before loading equivalent volumes.

12% Resolving gels and 4% stacking gels were prepared per Bio-Rad manuals, and SDS-PAGE performed on ice using the Mini-PROTEAN® system (Bio-Rad). Gels were run in standard Tris-glycine-SDS running buffer at 60 V through the stacking gel, then 100 V until resolved, alongside PageRuler™ Plus Prestained Protein Ladder (Thermo Fisher Scientific). Proteins were transferred to Immobilon®-PSQ PVDF membrane (Merck) using a Mini Trans-Blot system (Bio-Rad) on ice in Tris-glycine transfer buffer at 100 V for 20-60 min, depending on protein size. Membranes were cut horizontally where multiple targets were probed, washed in TBS-Tween, and blocked in 3% (w/v) skimmed milk in TBS-Tween (1 h, RT). Primary antibodies or antisera (in 3% milk/TBS-Tween) were incubated overnight at 4 °C, and membranes washed in TBS-Tween before secondary antibody (in 3% milk/TBS-Tween, 1 h, RT) and identical washing. Bands were revealed with SuperSignal™ West Pico PLUS substrate (Thermo Fisher Scientific) on a Syngene™ G:BOX Chemi XRQ. Antibodies are listed in Supplementary Table 1; antisera against RV-A16 3C was a gift from Sebastian Johnston and Roberto Solari (Imperial College London)^47^.

Where indicated, band intensities were quantified by densitometry in ImageJ (NIH), normalised to the β-Actin loading control, and expressed relative to the control condition.

### RT-qPCR

At the indicated time points, HeLa-H1 cells were washed in PBS and total RNA extracted using the High Pure RNA Isolation Kit (Roche) per the manufacturer’s instructions, except for the lysis step: 400 µl Lysis/Binding Buffer was diluted in 200 µl D-PBS and 600 µl added per well. Cells were scraped with an RNase-free tip, the whole-cell lysate collected and vortexed, transferred to a High Pure Filter Tube, and processed per instructions. Eluted RNA was quantified by NanoDrop™ One (RNA setting) and stored at −80 °C.

Purified RNA was reverse-transcribed using the High-Capacity cDNA RT Kit (Thermo Fisher Scientific): for each sample, RNA containing 250 ng was made up to 10 µl with RNase-free water. A 2× RT master mix containing random primers (10× RT buffer, 25× dNTP mix, 10× random primers, RT enzyme) was prepared per the manufacturer’s instructions for random-primed cDNA (RV-A16 viral RNA and 18S rRNA). Here, ‘viral RNA’ refers to total RV-A16 RNA, the vast majority of which during infection is positive-strand^48,49^. For all master mixes, a 2× no-RT control was prepared with the RT enzyme replaced by water. Master mix (10 µl) was added per 10 µl RNA (1× final, 20 µl total) and RT performed on a ProFlex PCR System (Thermo Fisher Scientific); cycling conditions are given in Supplementary Table 3. cDNA was stored at −20 °C.

Quantitative PCR reaction mixes were prepared using Luna® Universal Probe qPCR Master Mix (New England Biolabs) per the manufacturer’s instructions. Primer and probe concentrations are given in Supplementary Table 1. Mixes were aliquoted into white, opaque PCR plates (Sarstedt) and 12.5 ng cDNA added per well, with a no-RT control per gene per plate. For RV-A16 viral RNA, a standard curve was generated from ten DNA standards (pCR2_1_RV-A16, a gift from Sebastian Johnston, Imperial College London; serially diluted from 1×10^10^ to 1×10^1^ copies/µl)^42^. Samples were run in duplicate, and standards and no-RT controls in singlet, on a QuantStudio 1 (Thermo Fisher Scientific); cycling conditions are given in Supplementary Table 4.

For RV-A16 viral RNA, copy numbers per replicate were extrapolated from the standard curve, averaged, normalised to 18S rRNA, and multiplied by 4 (cDNA was made from 250 ng RNA) to give copies/µg RNA.

### Plasmid construction

Plasmid DNA was transformed into TOP10 chemically competent *E. coli* (Thermo Fisher Scientific) and purified using Monarch® Plasmid DNA Miniprep (New England Biolabs) or PureLink™ HiPure Plasmid Maxiprep (Thermo Fisher Scientific) kits, with glycerol stocks prepared for storage. DNA was quantified by NanoDrop™ One and analysed by agarose gel electrophoresis where required.

Site-directed mutagenesis was performed by inverse PCR using the KOD Hot Start DNA Polymerase Kit (Merck): reactions contained 1× buffer, 1.5 mM MgSO4, 0.2 mM each dNTP, 0.3 µM each primer, 2 ng/µl template and 0.02 U/µl polymerase in 50 µl (primers in Supplementary Table 1; cycling conditions in Supplementary Table 5). Methylated template was removed by DpnI digestion (4 h, 37 °C), and products were 5′ phosphorylated (T4 PNK) and self-ligated (Quick Ligation™ Kit; 50 ng, 5 min, 25 °C; all New England Biolabs) before transformation. Clones were verified by sequencing (Eurofins) and maxiprepped. To generate the replication-defective replicon, the RV-A16 nanoluciferase replicon plasmid (pRVA-16-NL, in which the P1 region is replaced by the NanoLuc® reporter under the RV-A16 IRES; a gift from Stanley Lemon and Kevin McKnight)^50^, was used as template, and the YGDD motif mutated to YGAA, with the 3D region sequence-verified.

### *In vitro* transcription and RNA purification

For *in vitro* transcription, plasmids were linearised (10 µg DNA, 2 h at 37 °C; all enzymes New England Biolabs): pR16.11 with SacI-HF® (with heat-inactivation); and pRVA-16-NL-YGAA with BamHI-HF®. Linearised DNA was purified (QIAquick PCR Purification Kit, Qiagen) and confirmed on an agarose gel against an undigested control. In vitro transcription used the MEGAscript™ T7 Kit (Thermo Fisher Scientific) for 4 h at 37 °C, after which RNA was treated with TURBO DNase, purified using in-house magnetic SPRI beads^51^, quantified by NanoDrop™ One, and transfected into HeLa-H1 cells (see ‘Cell transfections’).

### Statistical analysis

Data were analysed using the appropriate test (two-tailed paired *t*-test, one-way ANOVA) and *post-hoc* test (Dunnett’s) as outlined in the figure legends, using Prism (Graphpad). *, *P* < 0.05; **, *P* < 0.01; ***, *P* < 0.001; ****, *P* < 0.0001; ns, not significant.

## 6. Results

### Hsp70 activity is essential at an early stage of the RV-A16 replication cycle

We first sought to assess whether Hsp70 activity is required for RV-A16 replication. To this end, we examined the effect of VER-155008, an adenosine analogue that acts as an ATP-competitive inhibitor of Hsp70, arresting it in a half-open conformation and preventing the allosteric coupling required for its activity^52^. Remarkably, when HeLa-H1 cells were infected with RV-A16 in the presence of VER-155008, the production of infectious virus was inhibited in a dose-dependent manner, with complete inhibition at 50-100 µM (Figure 1A) and only minor reductions (22.9-30.4%) in cell viability (Figure 1B). These results indicate that Hsp70 activity is required for RV-A16 replication.

**Figure 1.**
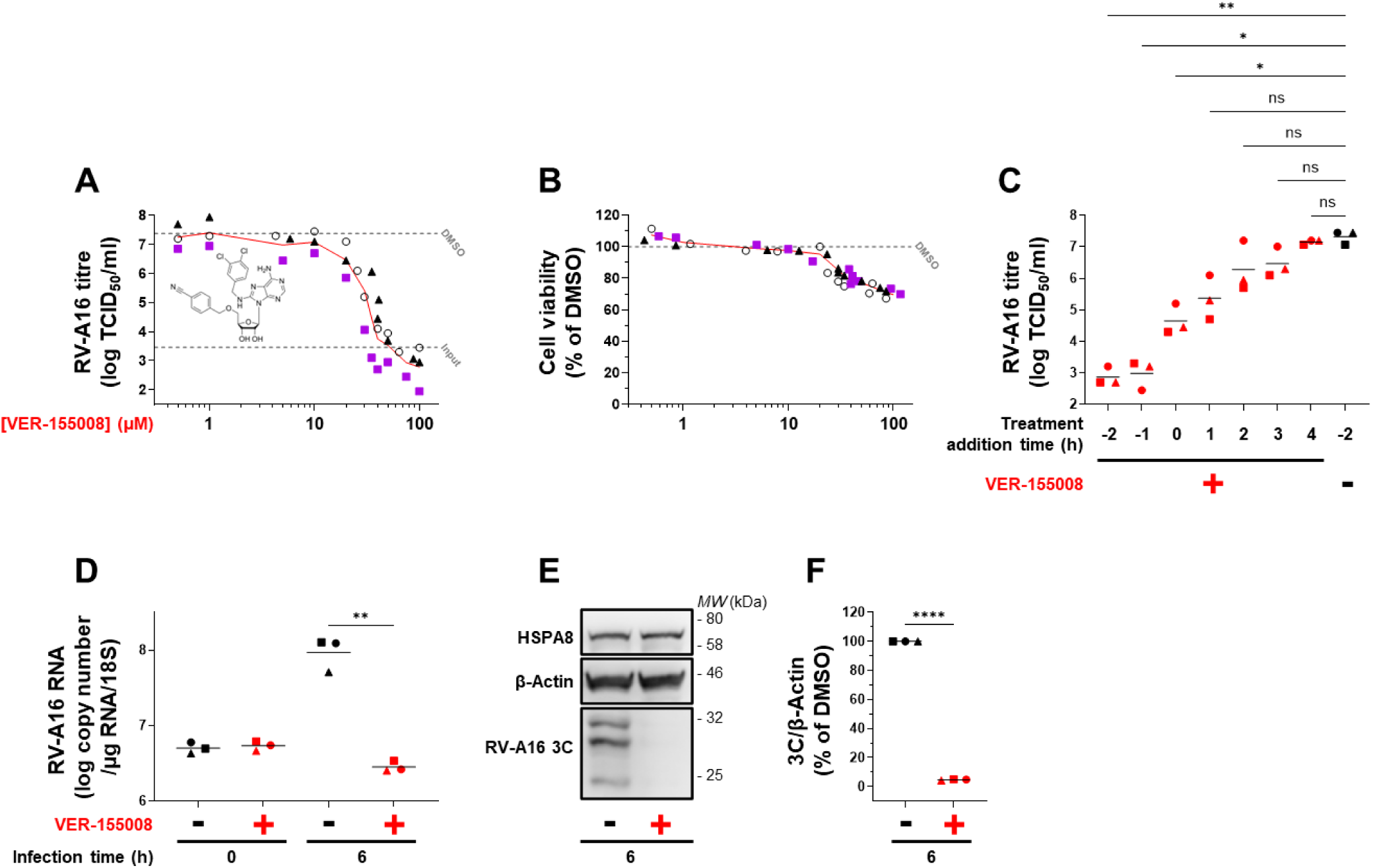
VER-155008, an ATP-competitive Hsp70 inhibitor, blocks an early stage of the RV-A16 replication cycle. **(A-B) VER-155008 blocks RV-A16 infectious virus production. (A)** Dose effect assay. HeLa-H1 cells were infected with RV-A16 (MOI 20) in the presence of DMSO or increasing concentrations of VER-155008 (0.5, 1, 5, 10, 20, 30, 35, 40, 50, 75 and 100 µM). Cells were re-treated with DMSO or the corresponding concentration of VER-155008 immediately after the virus adsorption. Viral titres were quantified at 0 hpi and 6 hpi by TCID_50_ assay (N=3). Viral titres in VER-155008-treated cells at 6 hpi are shown as individual points connected by a line. Drug structure is shown. Mean viral titres of DMSO-treated cells at 0 hpi (input) and 6 hpi are represented by dashed lines. **(B)** Viability assay. In parallel, HeLa-H1 cells were treated with DMSO or increasing concentrations of VER-155008 (as above), and cell viability was assessed at 7 h by resazurin assay (N=3). Data are presented as a percentage of the DMSO control. **(C)** VER-155008 time-of-addition assay. HeLa-H1 cells were infected with RV-A16 (MOI 20). Cells were treated with DMSO or 50 µM VER-155008 2 hours prior to the end of the virus adsorption (-2 h), at the start of the virus adsorption (-1 h), at the end of the virus adsorption (0 h) or at the indicated times post-infection. Viral titres were quantified at 6 hpi by TCID_50_ assay (N=3). (D-F) VER-155008 blocks RV-A16 RNA and NSP production. HeLa-H1 cells were infected with RV-A16 (MOI 20) in the presence of DMSO or 50 µM VER-155008. Cells were re-treated with DMSO or 50 µM VER-155008 immediately after the virus adsorption. (D) Viral RNA was quantified by RT-qPCR at 0 h and 6 h (N=3). (E) At 6 h, lysates were analysed by Western blotting for RV-A16 3C, HSPA8 and β-Actin (N=3). (F) 3C expression was quantified and normalised to β-Actin (N=3). For all graph panels (A-D, F), data are shown as individual points with means; where indicated, means are connected by lines. Points are coded by shape according to experimental replicate. Non-graph panel (E) shows a representative image. Statistical tests: (C) one-way ANOVA with Dunnett’s *post-hoc* test, (D, F) two-tailed paired t-test. *, *P* < 0.05; **, *P* < 0.01; ****, *P* < 0.0001; ns, not significant.

A time-of-addition experiment (Figure 1C), in which VER-155008 was added at defined times relative to the end of virus adsorption, showed that the full inhibitory effect of VER-155008 was obtained when added 1 or 2 hours pre-infection, but was progressively lost when added later. This suggests that Hsp70 activity is essential at an early stage of the viral replication cycle. Treatment of cells with VER-155008 also completely blocked the production of RV-A16 viral RNA (Figure 1D) and the 3C non-structural protein (NSP) (Figure 1E-F), further supporting a requirement for Hsp70 activity early in the viral replication cycle.

To confirm these findings using a mechanistically distinct inhibitor, we next examined the effect of pifithrin-µ (also named 2-phenylethynesulfonamide or PES) on RV-A16 replication. Unlike the ATP-competitive VER-155008, pifithrin-µ is an allosteric Hsp70 inhibitor that does not target the ATP-binding pocket^53^, but its precise mode of action remains incompletely understood^52^. When cells were pre-treated with pifithrin-µ and then infected with RV-A16 in the presence of the drug, the production of infectious virus was inhibited in a dose-dependent manner, with complete inhibition at 30 µM (Figure 2A) and only a minimal reduction (22.6%) in cell viability (Figure 2B). Like VER-155008, pifithrin-µ completely blocked the production of RV-A16 viral RNA (Figure 2C) and the 3C NSP (Figure 2D-E). The concordant effects of two mechanistically distinct Hsp70 inhibitors provide strong evidence that Hsp70 activity is essential during the early stages of RV-A16 replication.

**Figure 2.**
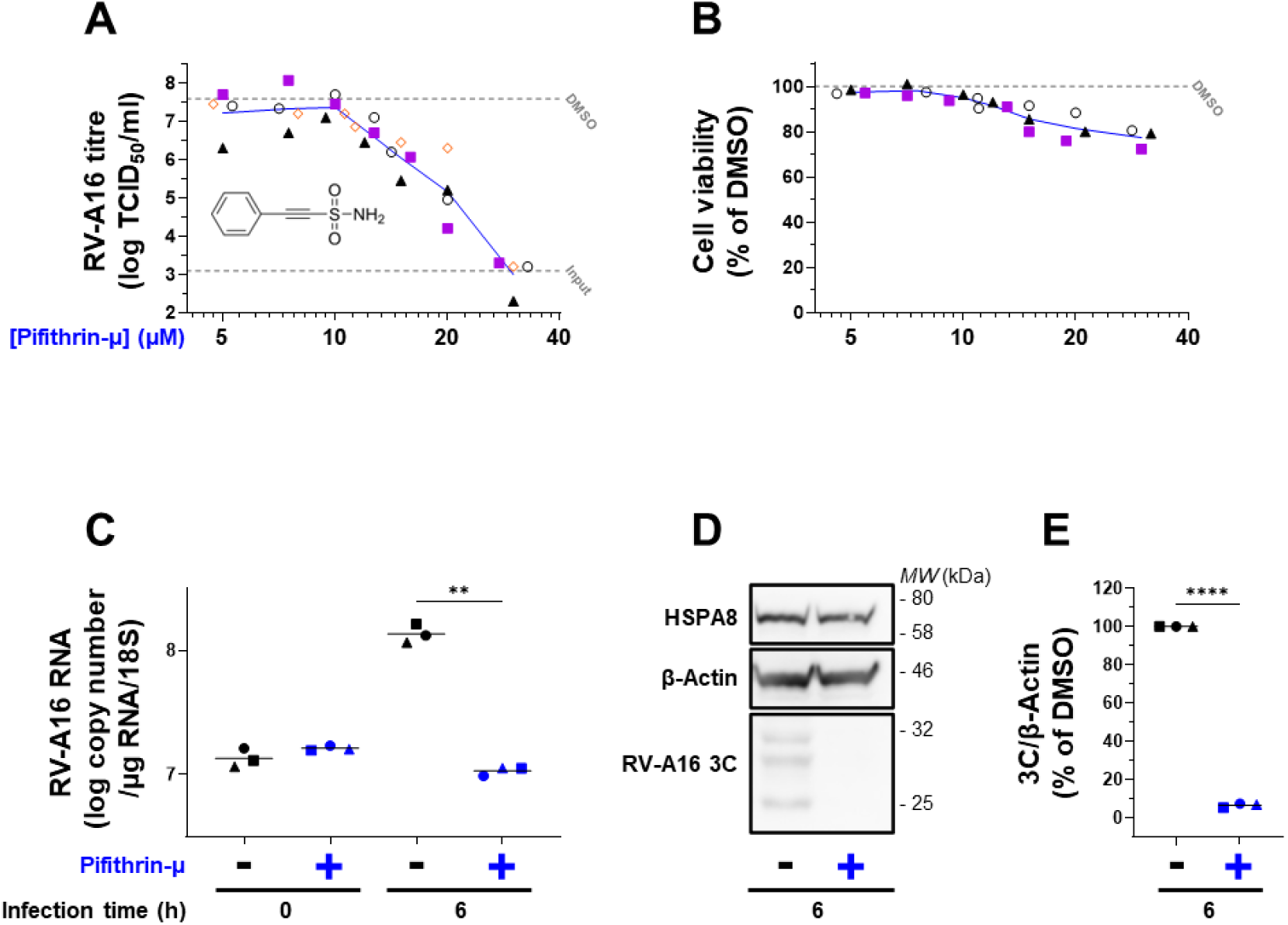
Pifithrin-µ, an allosteric Hsp70 inhibitor, blocks an early stage of the RV-A16 replication cycle. **(A-B) Pifithrin-µ blocks RV-A16 infectious virus production. (A)** Dose effect assay. HeLa-H1 cells were pre-treated for 1 h with DMSO or increasing concentrations of pifithrin-µ (5, 7.5, 10, 12, 15, 20 and 30 µM) and subsequently infected with RV-A16 (MOI 20) in the presence of DMSO or the corresponding concentration of pifithrin-µ. Cells were re-treated with DMSO or the corresponding concentration of pifithrin-µ immediately after the virus adsorption. Viral titres were quantified at 0 hpi and 6 hpi by TCID_50_ assay (N=4). Viral titres in pifithrin-µ-treated cells at 6 hpi are shown as individual points connected by a line. Drug structure is shown. Mean viral titres of DMSO-treated cells at 0 hpi (input) and 6 hpi are represented by dashed lines. **(B) Viability assay**. In parallel, HeLa-H1 cells were treated with DMSO or increasing concentrations of pifithrin-µ (as above), and cell viability was assessed at 8 h by resazurin assay (N=3). Data are presented as a percentage of the DMSO control. **(C-E) Pifithrin-µ blocks RV-A16 RNA and NSP production.** HeLa-H1 cells were pre-treated for 1 h with DMSO or 30 µM pifithrin-µ and subsequently infected with RV-A16 (MOI 20) in the presence of DMSO or 30 µM pifithrin-µ. Cells were re-treated with DMSO or 30 µM pifithrin-µ immediately after the virus adsorption. **(C)** Viral RNA was quantified by RT-qPCR at 0 h and 6 h (N=3). **(D)** At 6 h, lysates were analysed by Western blotting for RV-A16 3C, HSPA8 and β-Actin (N=3). **(E)** 3C expression was quantified and normalised to β-Actin (N=3). For all graph panels **(A-C, E)**, data are shown as individual points with means; where indicated, means are connected by lines. Points are coded by shape according to experimental replicate. Non-graph panel **(D)** shows a representative image. Statistical tests: **(C, E)** two-tailed paired t-test. **, *P* < 0.01; ****, *P* < 0.0001; ns, not significant.

### HSPA8, the major constitutively expressed Hsp70 family chaperone, is involved at an early stage of the RV-A16 replication cycle

To complement our pharmacological findings, we next assessed the role of HSPA8 (also named Hsc70) in RV-A16 replication. HSPA8 is the major constitutively expressed Hsp70 isoform that performs the bulk of housekeeping chaperone activity under non-stress conditions. Because HSPA8 is constitutively expressed, it is already present at the onset of infection, unlike inducible Hsp70 isoforms whose expression increases in response to cellular stress. Moreover, as HSPA8 is localised in the cytosol, where enterovirus translation and RNA replication occur^35^, it is well positioned to support these early stages of the viral replication cycle. siRNA-mediated knockdown reduced HSPA8 expression by 95% (Figure 3C, E) and resulted in a modest but significant reduction in the production of RV-A16 infectious virus (Figure 3A), viral RNA (Figure 3B), and 3C NSP (Figure 3C-D). These findings indicate that HSPA8 is involved in RV-A16 replication and are consistent with the conclusion from our inhibitor experiments that Hsp70 activity is required early in the viral replication cycle. Interestingly, siRNA knockdown of the JDPs DNAJA1 (HDJ2/DJ2) and DNAJC7 (DJ11/TPR2), which function as co-chaperones of the Hsp70 family, significantly reduced the production of RV-A16 infectious virus (Figure 3F) without significant reductions in cell viability (Figure 3G), suggesting that these factors are also involved in RV replication.

**Figure 3.**
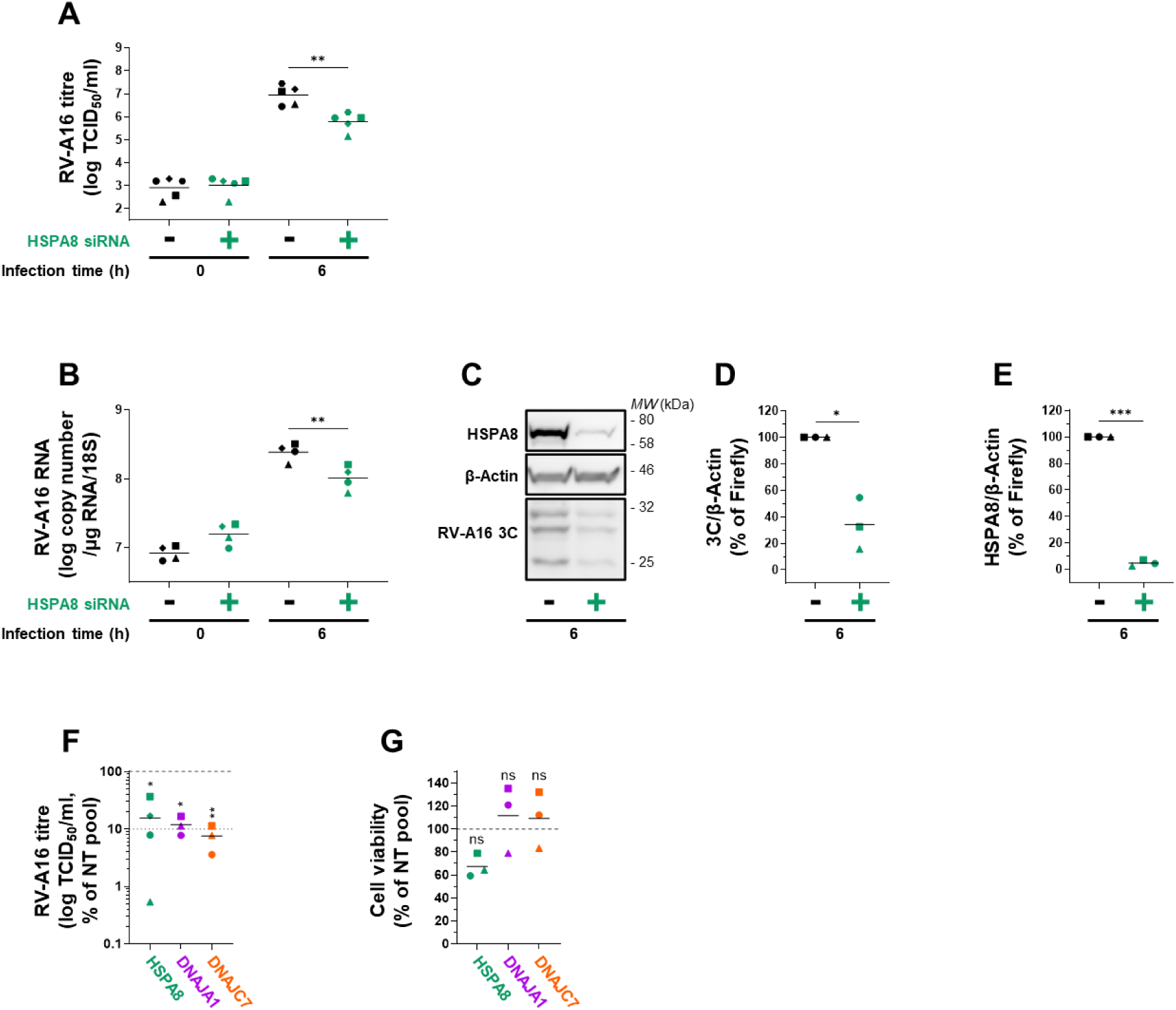
HSPA8, the major constitutively expressed Hsp70 family member, is involved at an early stage of the RV-A16 replication cycle. **(A-E) siRNA knockdown of HSPA8 inhibits the production of RV-A16 infectious virus, RNA, and NSP.** HeLa-H1 cells were transfected with 10 nM siRNA targeting HSPA8 or firefly luciferase for 72 h and then infected with RV-A16 (MOI 20). (A) Viral titres were quantified by TCID_50_ assay at 0 hpi and 6 hpi (N=5). **(B)** Viral RNA was quantified by RT-qPCR at 0 hpi and 6 hpi (N=4). (C) At 6 h, lysates were analysed by Western blotting for RV-A16 3C, HSPA8, and β-Actin (N=3). **(D-E)** Quantification of 3C and HSPA8 expression, normalised to β-Actin (N=3). **(F-G) siRNA knockdown of two Hsp70 co-chaperones, DNAJA1 and DNAJC7, inhibits the production of RV-A16 infectious virus. (F)** HeLa-H1 cells were transfected with 40 nM siRNA pools (X4 siRNA per pool) targeting HSPA8, DNAJA1 or DNAJC7, or with a non-targeting (NT) control siRNA pool, for 72 h. Transfected HeLa-H1 cells were infected with RV-A16 (MOI 20), and viral titres were quantified at 6 hpi by TCID_50_ assay (N=3-4). Data are presented as a percentage of the mean NT control (dashed grey line). 10% of the mean NT control is represented by a dotted grey line. (G) HeLa-H1 cells were transfected as above with siRNA pools targeting HSPA8, DNAJA1 or DNAJC7, or with an NT control siRNA pool. Cell viability was assessed at 72 h by resazurin assay (N=3). Data are presented as a percentage of the mean NT control (dashed grey line) For all graph panels **(A-B, D-G)**, data are shown as individual points with means. Points are coded by shape according to experimental replicate. Non-graph panel **(C)** shows a representative image. Statistical tests: **(A-B, D-E)** two-tailed paired t-test, (F-G) one-way ANOVA with Dunnett’s *post-hoc* test, comparing to the NT control pool. *, P < 0.05; **, *P* < 0.01; ***, *P* < 0.001; ns, not significant.

### Hsp70 activity is essential for IRES-mediated translation of RV RNA

As our infection experiments suggested an essential role for Hsp70 family members early in the viral replication cycle, we next sought to determine whether Hsp70 activity is required after viral entry. To this end, we bypassed normal viral entry by directly transfecting cells with RV-A16 RNA, in the presence or absence of VER-155008. In these conditions, VER-155008 treatment still completely abolished the production of infectious virus (Figure 4A), indicating that Hsp70 activity is indispensable for viral replication at a post-entry step.

**Figure 4.**
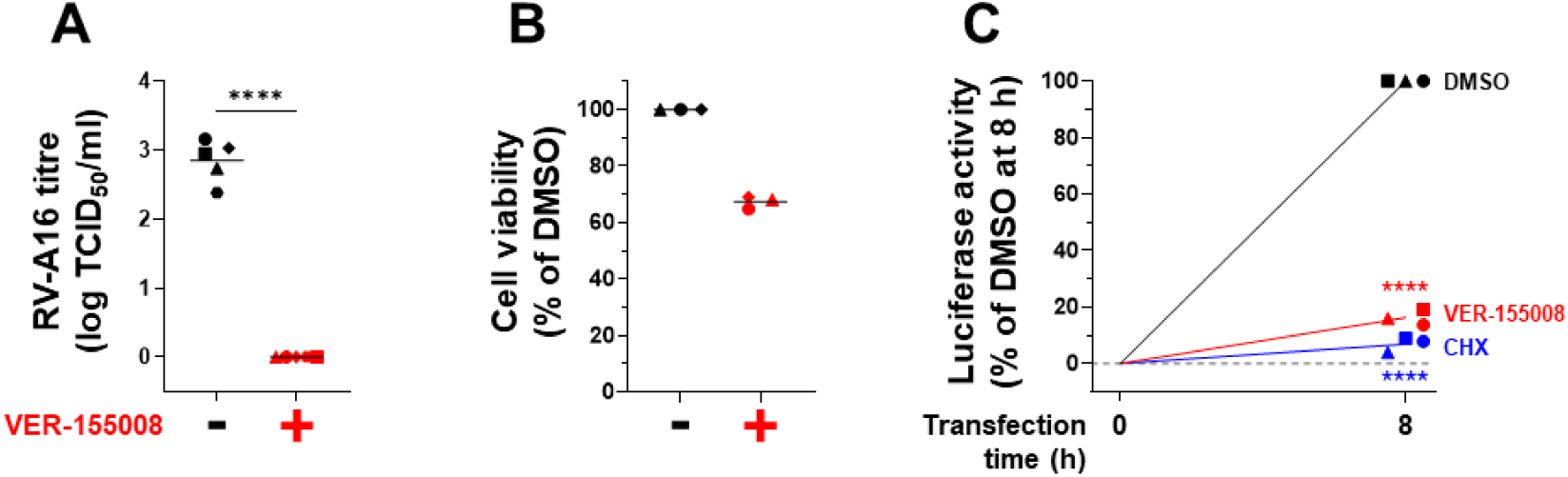
Hsp70 activity is essential for IRES-mediated translation of the incoming RV-A16 RNA. **(A-B) Hsp70 activity is required for a step occurring after RV-A16 entry. (A)** HeLa-H1 cells were pre-treated with DMSO or 50 µM VER-155008 for 1 h, and were subsequently transfected with 500 ng RV-A16 RNA in the presence of DMSO or 50 µM VER-155008. Viral titres were quantified at 8 h post-transfection by TCID_50_ assay (N=5). **(B)** In parallel, HeLa-H1 cells were treated with DMSO or 50 µM VER-155008 in the absence of transfection. Cell viability was assessed at 9 h post-treatment by resazurin assay (N=3). **(C) Hsp70 activity is essential for RV-A16 IRES-mediated translation.** HeLa-H1 cells were pre-treated with DMSO, 50 µM VER-155008 or 178 µM cycloheximide (CHX) for 1 h, and were subsequently transfected with 100 ng replication-defective RV-A16 nanoluciferase replicon (with the conserved 3D polymerase active-site motif YGDD mutated to YGAA) in the presence of DMSO, 50 µM VER-155008, or 178 µM CHX. Luciferase activity was measured at the indicated times. Values were t=0-subtracted and normalised to the DMSO 8 h post-transfection value within each experiment (N=3). Means are connected by lines; the 0 h baseline is shown as a dashed grey line. Data are shown as individual points with means; where indicated, means are connected by lines. Points are coded by shape according to experimental replicate. Statistical tests: **(A)** two-tailed paired t-test, **(C)** two-way ANOVA, comparing drug treatments to the DMSO control at each time point. ****, *P* < 0.0001; ns, not significant.

As is the case for related picornaviruses, RV RNA contains a highly structured RNA element in the 5’ untranslated region (UTR) known as an internal ribosome entry site (IRES), which permits cap-independent translation of the viral RNA by host ribosomes in conjunction with numerous canonical and IRES *trans*-acting factors (ITAFs). Following entry into the cell cytosol, the viral RNA is translated into a single polyprotein, which is subsequently cleaved by virus-encoded proteases to yield individual structural and non-structural proteins. The viral RNA is then replicated, and new copies can be used for further rounds of translation and replication. Because newly synthesized viral non-structural proteins only become detectable by western blot and immunofluorescence after 4 hours of infection, once the viral RNA has been sufficiently amplified through replication^54^, reduced detection of 3C after Hsp70 inhibition (Figure 1E-F, Figure 2D-E) or HSPA8 knockdown (Figure 3C-D) could reflect impaired RNA replication rather than a direct effect on IRES-mediated translation. To specifically assess the effect of VER-155008 on IRES-mediated translation, we transfected cells with a non-replicative luciferase reporter RNA, in which luciferase expression is under the control of the RV-A16 IRES. Remarkably, VER-155008 inhibited the expression of the luciferase reporter to a level comparable to that of cycloheximide, a protein translation inhibitor (Figure 4C), demonstrating that Hsp70 activity is essential for RV-A16 IRES-mediated translation.

## 7. Discussion

In this study, we identify Hsp70 family chaperones as essential host factors for RV-A16 replication and demonstrate that Hsp70 activity is required at an early post-entry stage of the viral replication cycle. Using two mechanistically distinct Hsp70 inhibitors, together with siRNA-mediated knockdown of the constitutively expressed Hsp70 isoform HSPA8 and the co-chaperones DNAJA1 and DNAJC7, we show that disruption of the Hsp70 chaperone system impairs RV-A16 replication. Our data indicate that Hsp70 activity is required early in the viral replication cycle, as Hsp70 inhibitors prevented the accumulation of viral RNA and the 3C non-structural protein and were most effective when present during the initial stages of infection. However, Hsp70 inhibition remained effective when viral entry was bypassed by direct transfection of the viral RNA, demonstrating that its antiviral effect occurs after entry. Importantly, using a specific translation reporter, we show that Hsp70 activity is required for RV-A16 IRES-mediated translation. Together, these findings provide the first direct evidence that RVs rely on Hsp70 chaperones for replication and identify viral translation as a key Hsp70-dependent step in the RV replication cycle.

The requirement for Hsp70 activity in RV-A16 IRES-mediated translation is consistent with previous studies of related enteroviruses. Pharmacological inhibition of Hsp70 activity with JG40 significantly reduced luciferase expression from EV-A71 IRES translation reporters^31^, while knockdown or knockout of the Hsp70 isoforms HSPA1A/HSPA1B, HSPA6, HSPA8 or HSPA9 also impaired reporter activity^31,32^, supporting a role for Hsp70 chaperones in EV-A71 translation. Together with our findings on RV-A16, these observations suggest that Hsp70-dependent viral translation may be a conserved feature of the enterovirus replication cycle. However, the precise mechanisms by which Hsp70 chaperones promote viral translation have yet to be defined.

IRES-mediated translation in enteroviruses relies on a range of canonical cellular translation factors, including eIF4A, eIF4B, eIF3, eIF2 and eIF1A^55^, as well as a diverse group of IRES *trans*-acting factors (ITAFs) that bind the IRES and promote recruitment of the translation machinery. ITAFs implicated in the translation of RV, PV, or EV-A71 include poly(rC)-binding proteins 1 and 2 (PCBP1/2)^56–60^, polypyrimidine tract binding protein 1 (PTBP1)^61–63^, CSDE1 (also known as unr, upstream of N-ras)^64,65^, lupus autoantigen (La)^66,67^, heterogeneous nuclear ribonucleoprotein A1 (hnRNP A1)^68,69^, HuR, Ago2^70^, the 68-kDa Src-associated protein in mitosis (Sam68)^71^, far-upstream element-binding protein 1 (FBP1)^72^, and the DDX3X RNA helicase^73^. Of these, PCBP2 is particularly important, binding the stem-loop IV of the IRES to form a ribonucleoprotein (RNP) complex that promotes translation initiation^56,57^. Following translation initiation, the viral protease 2A^pro^ cleaves several host factors, including eIF4GI, eIF4GII and polyadenylate-binding protein 1 (PABP1), to shut off cap-dependent translation of host mRNAs and favour viral protein synthesis^74–76^. Subsequently, the C-terminal fragment of cleaved eIF4G binds the IRES to further enhance translation^77^. Given the diverse roles of Hsp70 chaperones in protein folding, protein complex assembly and the regulation of protein activity, it is conceivable that they could facilitate the function of one or more of these host factors, or their assembly into complexes required for efficient IRES-mediated translation. Moreover, as Hsp70 activity is directed by specific co-chaperones, distinct Hsp70-JDP complexes may regulate different stages of the viral translation process.

Our siRNA knockdown experiments provide further insight into the contribution of Hsp70 network components to RV replication. Compared with pharmacological inhibition using VER-155008 or pifithrin-µ, which broadly inhibit Hsp70 family activity and completely abolish RV replication, knockdown of HSPA8 alone resulted in a modest but significant reduction in the production of RV RNA, 3C non-structural protein and infectious viral particles. Several explanations may account for the difference in the magnitude of these effects. First, additional Hsp70 isoforms may contribute to RV replication, as reported for EV-A71^26,31,32^. Second, functional redundancy within the Hsp70 family^35^ may allow other isoforms to compensate, at least in part, when HSPA8 expression is reduced. Finally, although the siRNA reduced HSPA8 expression by around 95%, residual HSPA8 may have been sufficient to support low levels of viral replication. Future work could directly test the contribution of other Hsp70 isoforms, particularly HSPA1A/1B, HSPA6 and HSPA9, which have been implicated in EV-A71 replication^26,31,32^, and determine whether combined knockdown of multiple isoforms can reproduce the complete inhibition of viral replication observed with VER-155008 and pifithrin-µ. Nonetheless, our data indicate a role for HSPA8 in the RV replication cycle, acting prior to, or at the onset of viral RNA replication. As HSPA8 is constitutively expressed in the cytosol under basal conditions^35^, it is well positioned to support early stages of infection prior to the induction of stress-inducible Hsp70 isoforms. Together with our inhibitor data, these findings raise the possibility that HSPA8 contributes to the initiation and/or enhancement of RV IRES-mediated translation, possibly in conjunction with other Hsp70 isoforms, viral proteins or ITAFs. This would be consistent with studies on EV-A71, for which HSPA8 was shown to be required for IRES-mediated translation of the viral RNA^26^.

In line with the established requirement for co-chaperones in Hsp70 function^35^, siRNA-mediated depletion of the JDPs DNAJA1 and DNAJC7 also impaired RV replication, indicating that these co-chaperones may support Hsp70-dependent processes during RV infection, potentially in concert with HSPA8 during the early stages of the viral replication cycle. A contribution of Hsp70 chaperones and their JDP co-chaperones to viral replication has precedent among positive-strand RNA viruses beyond picornaviruses, particularly among flaviviruses. In dengue and Zika virus infection, pharmacological inhibition of Hsp70 activity or knockdown of HSPA8 impairs viral replication^29,30^. In addition, several JDP co-chaperones have been implicated in different stages of the dengue virus replication cycle^29^, supporting the broader concept that viral replication can depend on specific components of the Hsp70 co-chaperone network, in which specific JDPs can direct Hsp70 chaperone activity to defined steps of the viral replication cycle. Whether DNAJA1 and DNAJC7 act together with HSPA8 during RV-A16 replication, and whether they contribute directly to IRES-mediated translation or to another step of the viral replication cycle, remains to be determined.

The identification of Hsp70 chaperones as essential host factors for RV replication may have important implications for antiviral development. Host-targeted antiviral strategies directed against conserved cellular pathways offer the potential for broad-spectrum activity and, compared with direct-acting antivirals, may present a higher barrier to the emergence of viral resistance^78–80^. In this context, our findings further support the notion that Hsp70 chaperones act as “pan-enterovirus” host factors, extending the evidence from EV-A71, CVA16, CVB1, CVB3 and E11 to include RV-A16. Encouragingly, Hsp70-targeted strategies have shown promise beyond enteroviruses. Indeed, Hsp70 inhibitors suppress replication of multiple flaviviruses in disease-relevant human cell models and protect mice from lethal Zika virus infection, while reducing viraemia and disease severity without overt host toxicity^29,30^. Moreover, serial passaging of dengue and Zika virus in the presence of Hsp70 inhibitors did not readily select for resistant viruses^29,30^, supporting the idea that Hsp70 inhibition may present a high barrier to viral escape. Importantly, although inhibition of such central chaperones raises potential concerns about disruption of host cell proteostasis, allosteric inhibitors of the Hsp70-NEF complex have been reported to show limited toxicity at concentrations that inhibit viral replication, likely because they leave other chaperone functions intact^29,30^. Viral infection may also create a heightened dependence on specific Hsp70-mediated processes compared with basal host cell proteostasis^29,30^. Together, these findings suggest that therapeutically useful antiviral windows may be achievable. Future work could further optimise Hsp70-targeted approaches for potency, selectivity and tolerability, for example by targeting virus-specific Hsp70 functional complexes, including interactions between Hsp70 isoforms, co-chaperones and viral or cellular clients.

In summary, we demonstrate that Hsp70 activity is indispensable for the replication of RV-A16 and define a critical role for these proteins in IRES-mediated translation of the viral RNA. These findings expand our understanding of the host factor network exploited by RVs, further establish Hsp70 chaperones as important “pan-enterovirus” host factors, and highlight the potential of targeting Hsp70-associated pathways to control RV infection.

## 10. Author contributions

Conceptualization, A.M., M.T.J.; Data Curation, M.T.J., A.O.M., A.M.; Formal Analysis, M.T.J., A.M.; Funding Acquisition, A.M.; Investigation, M.T.J., C.D., A.O.M.; Methodology, M.T.J., C.D., A.M.; Project Administration, A.M.; Resources, A.M.; Supervision, A.M.; Visualization, M.T.J.; Writing - original draft, M.T.J.; Writing - review & editing, M.T.J., C.D., A.O.M. and A.M.

## 11. Acknowledgements

We thank Sebastian Johnston and Roberto Solari for the gift of plasmid pCR2_1_RV-A16 and antibodies against RV-A16 non-structural protein 3C. We thank Stanley Lemon and Kevin McKnight for the gift of plasmids pR16.11 and pRVA-16-NL.

