## Supplementary Tables for "The heat shock protein 70 (Hsp70) family of chaperones is essential for rhinovirus replication"

| REAGENT or RESOURCE | SOURCE | IDENTIFIER |
|---|---|---|
| **Antibodies** | | |
| Goat polyclonal anti-mouse IgG (H+L), HRP-conjugated | Jackson ImmunoResearch | 115-035-146 |
| Goat polyclonal anti-rabbit IgG (H+L), HRP-conjugated | Jackson ImmunoResearch | 111-035-144 |
| Mouse monoclonal anti-β-Actin | Proteintech | 60008-1-Ig |
| Mouse monoclonal anti-HSPA8 | Santa Cruz Biotechnology | sc-7298 |
| Rabbit polyclonal anti-RV-A16 3C | Gifted by Sebastian Johnston and Roberto Solari[47] | N/A |
| **Bacterial and virus strains** | | |
| One Shot™ TOP10 Chemically Competent E. coli | ThermoFisher Scientific | C404003 |
| RV-A16 | ATCC | VR-283 |
| **Buffers and media** | | |
| 2X Laemmli sample buffer: 0.15 M Tris-HCl pH 6.8, 25% (v/v) glycerol, 5% (w/v) SDS, RNase-free water, supplemented with 0.2 M DTT | Made in-house | N/A |
| 5X siRNA buffer | Dharmacon | B-002000-UB-100 |
| Complete medium: high glucose Dulbecco's modified Eagle's medium supplemented with 10% (v/v) fetal bovine serum | Made in-house | N/A |
| Enzyme buffer: rCutSmart® Buffer 10X | New England Biolabs | B6004S |
| Enzyme buffer: T4 Polynucleotide Kinase Reaction Buffer 10X | New England Biolabs | B0201S |
| Infection medium: high glucose Dulbecco's modified Eagle's medium supplemented with 2% (v/v) fetal bovine serum and 25 mM HEPES (pH 7.2-7.5) | Made in-house | N/A |
| PBS-Tween: 0.05% (v/v) Tween-20 in PBS | Made in-house | N/A |
| SDS-PAGE running buffer: 0.19 M glycine, 25 mM Tris, 0.1% (w/v) SDS | Made in-house | N/A |
| siMAX Universal Buffer 5X | Eurofins | N/A |
| TBS-Tween: 20 mM Tris, 150 mM NaCl, pH 7.4–7.6, 0.1% (v/v) Tween-20 | Made in-house | N/A |
| Titration medium: high glucose Dulbecco's modified Eagle's medium supplemented with 2% (v/v) fetal bovine serum, 25 mM HEPES (pH 7.2-7.5) and 1% (v/v) penicillin/streptomycin | Made in-house | N/A |
| Transfer buffer: 0.19 M glycine, 25 mM Tris | Made in-house | N/A |
| Tris-acetate-EDTA (TAE) buffer 50X | Thermo Fisher Scientific | BP1332-1 |
| **Chemicals and reagents** | | |
| 37% hydrochloric acid | VWR | 20252.295 |
| 5X siRNA buffer | Dharmacon | B-002000-UB-100 |
| Acrylamide/bis-acrylamide | Merck | A3449-100ML |
| Agarose powder | Thermo Fisher Scientific | 16500-500 |
| Ammonium persulfate (APS) | Thermo Fisher Scientific | 17874 |
| Ampicillin sodium salt | Merck | A9518-25G |
| ATP sodium solution (100mM) | RayBiotech | 366-10001-1 |
| Bromophenol blue | Alfa Aesar | 32641 |
| Cycloheximide (CHX) | Thermo Fisher Scientific | J66665.03 |
| Dimethyl sulfoxide (DMSO) | Merck | D8418 |
| Dithiothreitol (DTT) | Fluorochem | M02712 |
| Enzyme buffer: rCutSmart® Buffer 10X | New England Biolabs | B6004S |
| Enzyme buffer: T4 Polynucleotide Kinase Reaction Buffer 10X | New England Biolabs | B0201S |
| Enzyme: BamHI-HF® (20,000 units/ml) | New England Biolabs | R3136S |
| Enzyme: DpnI (20,000 units/ml) | New England Biolabs | R0176 |
| Enzyme: SacI-HF® (20,000 units/ml) | New England Biolabs | R3156S |
| Enzyme: T4 Polynucleotide Kinase (10,000 units/ml) | New England Biolabs | M0201S |
| Fetal bovine serum (FBS) | Thermo Fisher Scientific | A5256701 |
| Gel Loading Dye, Purple (6X), no SDS | New England Biolabs | B7025S |
| Glycerol | Thermo Fisher Scientific | A16205.AP |
| Glycine | VWR | 10119CU |
| HEPES | Merck | H3375-100G |
| High glucose Dulbecco's modified Eagle's medium (DMEM) | Thermo Fisher Scientific | 41965039 |
| LB agar, Vegitone | Merck | 19344-500G-F |
| LB broth, high salt | Merck | 51208-500G-F |
| Lipofectamine® RNAiMAX transfection reagent | Thermo Fisher Scientific | 13778 |
| Midori Green Direct DNA Stain | Geneflow | S6-0016 |
| Molecular-grade water | Merck | 12190000 |

| | | |
|---|---|---|
| mRNA Boost reagent | Mirus Bio | MIR 2255 |
| Opti-MEM® I reduced serum medium | Thermo Fisher Scientific | 31985-047 |
| PageRuler™ Plus Prestained Protein Ladder, 10 to 250 kDa | Thermo Fisher Scientific | 26619 |
| Penicillin-streptomycin (10,000 U/mL) | Thermo Fisher Scientific | 15140122 |
| Pifithrin-μ | Merck | P0122-5MG |
| Quick-Load® 1 kb Plus DNA Ladder | New England Biolabs | N0469S |
| Resazurin powder | Fluorochem | F492502 |
| Rnase free water | Merck | W4502-1L |
| siMAX Universal Buffer 5X | Eurofins | N/A |
| Skimmed milk | Merck | 70166-500G |
| Sodium chloride (NaCl) | Thermo Fisher Scientific | S/3160/65 |
| Sodium dodecyl sulfate (SDS) | Merck | 75746-250G |
| SPRI RNA purification beads | Made in-house[51] | N/A |
| TBS-Tween: 20 mM Tris, 150 mM NaCl, pH 7.4–7.6, 0.1% (v/v) Tween-20 | Made in-house | N/A |
| Tetramethylethylenediamine (TEMED) | Merck | T9281-25ML |
| TransIT-mRNA reagent | Mirus Bio | MIR 2255 |
| Trizma® base | Merck | T6066-1KG |
| Trypsin-EDTA (0.25%), phenol red | Thermo Fisher Scientific | 25200072 |
| Tween-20 | Merck | P7949-500ML |
| VER-155008 | Neo Biotech | NB-64-38120-5mg |
| **Commercial assays** | | |
| Applied Biosystems™ High-Capacity cDNA Reverse Transcription (RT) Kit | Thermo Fisher Scientific | 4368814 |
| High Pure RNA Isolation Kit | Roche | 11828665001 |
| Luna® Universal Probe qPCR Master Mix | New England Biolabs | M3004X |
| MEGAscript™ T7 Transcription Kit | Thermo Fisher Scientific | AMB13345 |
| Monarch® PCR & DNA Cleanup Kit | New England Biolabs | T1030L |
| Monarch® Plasmid DNA Miniprep Kit | New England Biolabs | T1010L |
| Nano-Glo® Luciferase Assay Kit | Promega | N1120 |
| Novagen® KOD Hot Start DNA Polymerase Kit | Merck | 71086-4 |
| PureLink™ HiPure Plasmid Maxiprep Kit | Thermo Fisher Scientific | K210007 |
| QIAquick PCR Purification Kit for PCR Cleanup | Qiagen | 28106 |
| Quick Ligation™ Kit | New England Biolabs | M2200L |
| SuperSignal™ West Pico PLUS Chemiluminescent Substrate | Thermo Fisher Scientific | 34580 |
| **Experimental models: cell lines** | | |
| Human: HeLa-H1 cells | ATCC | CRL-1958 |
| **Oligonucleotides** | | |
| RT-qPCR: 18S rRNA forward primer: 5'- CGCCGCTAGAGGTGAAATTCT -3' (working concentration 0.4 μM) | Merck | OLAM347 |
| RT-qPCR: 18S rRNA probe: 5'- [FAM]ACCGGCGCAAGACGGACCAGA[TAM] -3' (working concentration 0.2 μM) | Eurofins | N/A |
| RT-qPCR: 18S rRNA reverse primer: 5'- CATTCTTGGCAAATGCTTTCG -3' (working concentration 0.4 μM) | Merck | OLAM348 |
| RT-qPCR: RV-A16 viral RNA forward primer: 5'- GTGAAGAGCCGCGTGTGCT -3' (working concentration 0.05 μM) | Merck | OLAM345 |
| RT-qPCR: RV-A16 viral RNA probe: 5'- [FAM]TGAGTCCTCCGGCCCCTGAATG[TAM] -3' (working concentration 0.2 μM) | Eurofins | N/A |
| RT-qPCR: RV-A16 viral RNA reverse primer: 5'- GCTGCAGGTTTAAGGTTAGCC -3' (working concentration 0.3 μM) | Merck | OLAM346 |
| SDM: RVA-16-NL-YGAA forward primer: 5'- GcTGTGATCTTTTCTTACAAGTATAAACTAGAC -3' (working concentration 0.3 μM) | Merck | OLAM358 |
| SDM: RVA-16-NL-YGAA reverse primer: 5'- AgCACCATAGGCAATTATTTTAAGTTTATCTAA -3' (working concentration 0.3 μM) | Merck | OLAM359 |
| Single siRNA: firefly luciferase antisense: 5′- UCGAAGUAUUCCGCGUACGug -3′ | Eurofins | siAM1_Luc GL2_Ctl |
| Single siRNA: firefly luciferase sense: 5′- CGUACGCGGAAUACUUCGAtt -3′ | Eurofins | siAM1_Luc GL2_Ctl |
| Single siRNA: HSPA8 antisense: 5'- AGAAUAGGUAGUGAAGGUCtg -3' | Eurofins[45] | siAM9_Hsc70_1 |
| Single siRNA: HSPA8 sense: 5'- GACCUUCACUACCUAUUCUga -3' | Eurofins[45] | siAM9_Hsc70_1 |
| siRNA screen: HSPA8 cherry-pick ON-TARGETplus siRNA library SMARTpool | Dharmacon | L-017609-00 |
| siRNA screen: DNAJA1 cherry-pick ON-TARGETplus siRNA library SMARTpool | Dharmacon | L-019617-00 |
| siRNA screen: DNAJC7 cherry-pick ON-TARGETplus siRNA library SMARTpool | Dharmacon | L-019566-01 |

| Recombinant DNA | | |
|---|---|---|
| pCR2_1_RV-A16 (plasmid encoding an 878 bp fragment of RV-A16 genome, for use as DNA standard for absolute quantification of RV-A16 viral RNA by RT-qPCR) | Gifted by Sebastian Johnston | N/A |
| pR16.11 (full-length RV-A16) | Gifted by Stanley Lemon and Kevin McKnight[46] | N/A |
| RVA-16-NL (RV-A16 nanoluciferase replicon) | Gifted by Stanley Lemon and Kevin McKnight[50] | N/A |
| RVA-16-NL-YGAA (RV-A16 replication-defective nanoluciferase replicon) | This study | N/A |
| **Software and algorithms** | | |
| Prism (v10.5.0) | GraphPad | https://www.graphpad.com/ |
| **Other** | | |
| Applied Biosystems™ ProFlex PCR System | Thermo Fisher Scientific | 4484073 |
| Applied Biosystems™ QuantStudio 1 Real-Time PCR System | Thermo Fisher Scientific | A40426 |
| Black, opaque 96-well plates | Greiner Bio-One | 655986 |
| FLUOstar® Omega plate reader | BMG Labtech | 415-101-AFL |
| Immobilon® -PSQ PVDF Membrane 0.2 µM pore size | Merck | ISEQ00010 |
| Mini-PROTEAN® electrophoresis system | Bio-Rad | 1658004 |
| NanoDrop™ One microvolume UV-vis spectrophotometer | Thermo Fisher Scientific | ND-ONE-W |
| Optically transparent film | Sarstedt | 95.1999 |
| Syngene™ G:BOX Chemi XRQ image analyser | Thermo Fisher Scientific | 15472280 |

**Supplementary Table 1. List of key resources used in this study.**

| Well | Opti-MEM® I (µl) | RNA (µg) | *Trans*IT-mRNA reagent (µl) | mRNA Boost reagent (µl) | HeLa-H1 in complete medium (µl) | Total volume to seed (µl) |
|---|---|---|---|---|---|---|
| 96-well plate | 10 | 0.1 | 0.2 | 0.2 | 90 | 100 |
| 24-well plate | 50 | 0.5 | 0.5 | 0.5 | 450 | 500 |

**Supplementary Table 2. Reverse transfection of *in vitro* transcribed RNA.**

| Step | Temperature (°C) | Time (minutes) | Number of cycles |
|---|---|---|---|
| 1 | 25 | 10:00 | X1 |
| 2 | 37 | 120:00 | X1 |
| 3 | 85 | 5:00 | X1 |
| 4 | 4 | Hold | X1 |

**Supplementary Table 3. PCR protocol for RT of cDNA.**

| Step | Temperature | Time (seconds) | Number of cycles |
|---|---|---|---|
| 1 | 95 | 00:60 | X1 |
| 2 | 95 | 00:15 | X40 |
| 3 | 60 | 00:30 (+ plate read) | |

**Supplementary Table 4. RT-qPCR protocol.**

| Step | Temperature (°C) | Time (minutes) | Number of cycles |
|---|---|---|---|
| 1. Polymerase activation | 95 | 02:00 | X1 |
| 2. Denaturation | 95 | 00:20 | X40 |
| 3. Annealing | Annealing Temp | 00:10 | |
| 4. Extension | 70 | 00:25 per 1 kb | |
| 5. Final extension | 70 | 10:00 | X1 |
| 6. Hold | 4 | Hold | X1 |

**Supplementary Table 5. PCR protocol for site-directed mutagenesis.**
